# HMGB1 orchestrates uterine macrophage trafficking to safeguard embryo implantation

**DOI:** 10.1101/522490

**Authors:** Shizu Aikawa, Wenbo Deng, Xiaohuan Liang, Jia Yuan, Amanda Bartos, Xiaofei Sun, Sudhansu K Dey

## Abstract

A reciprocal communication between the implantation-competent blastocyst and the receptive uterus is essential to implantation. Blastocyst implantation is considered to be a regulated proinflammatory response in the uterus, however the underlining mechanism remains elusive. Here, we provide genetic evidence that High-mobility group protein Box-1 (HMGB1), expressed in uterine cell nuclei, restricts inflammatory responses during the periimplantation period. Conditional deletion of uterine *Hmgb1* by using a *Pgr-Cre* driver (*Pgr*^*cre*/+^*Hmgb1*^*f/f*^) shows substantial infertility because of defective implantation and subsequent adverse ripple effects. These mice accumulate and retain an increased number of macrophages in the stroma on day 4 of pregnancy with a unique enrichment of macrophages in the stroma encircling the blastocyst on day 5, evoking inflammatory responses. These results are in contrast to previous findings that HMBG1 is an internal alarmin. In search for the mechanism, we found that *Hmgb1*-deleted stromal cells show reduced activation of PR and decreased *Hoxa10* expression, providing evidence that PR and Hoxa10 mediated regression of inflammation is mediated by HMGB1. In addition, levels of two macrophage attractants CSF1 and CCL2 are elevated in the stroma and in vitro studies show that CSF1 specifically attracts macrophages which is abrogated if challenged with a CSF1 receptor antagonist. The results suggest that *Hmgb1* contributes to successful blastocyst implantation by regulating macrophage trafficking in the stroma to prevent excessive inflammatory responses.

---

An orchestrated reciprocal interaction between an implantation-competent blastocyst and a receptive uterus is critical for healthy implantation which sets the stages for subsequent pregnancy success^1,2^. Defects during implantation either terminate pregnancy, or compromise decidualization, placentation and parturition, finally impacting pregnancy outcomes^1,3,4,5^. In mice, the uterus becomes receptive to blastocyst implantation on day 4 of pregnancy (day 1 = vaginal plug), but undergoes refractory condition by day 5^1,2^. Blood vessels enter the uterus from the mesometrium with orientation of the uterus along a mesometrial-antimesometrial (M-AM) axis. On the evening of day 4, planar cell polarity (PCP) signaling allows luminal epithelial (LE) evaginations toward the AM pole to form a specialized crypt (implantation chamber) along with pre-existing glands, establishing a direct communication between the blastocyst and glands^6^. This process is accompanied by increased endometrial vascular permeability at the site of blastocyst apposition, which can be visualized as blue bands after an injection of blue dye solution^7^. Stromal cells begin to proliferate on day 3 and become more intense on day 4. On day 5 following implantation, stromal cells undergo proliferation and differentiation to decidual cells (decidualization) surrounding the blastocyst^1^.

After coitus, the mouse uterus is accompanied by immune cell infiltration with increased levels of inflammatory cytokines on days 1 and 2 of pregnancy. However, these conditions subside on day 3, the day before embryos enter the uterine cavity^8,9^. Macrophages progressively migrate away from the stroma with progression of pregnancy with the rise of progesterone (P_4_) levels from the newly formed corpora lutea, suggesting detrimental effects of macrophages on blastocyst survival and implantation. It remains unclear how the uterus quells immune cell infiltration and inflammatory signals to protect blastocyst implantation.

High Mobility Group Box Protein-1 (HMGB1) was originally identified as a nuclear protein bound to nucleosomes to maintain chromatin structure and gene transcriptions. HMGB1 is considered a damage associated molecular pattern molecule (DAMP), since under certain stress conditions or necrosis, HMGB1 is translocated from nucleus into cytosol or released into extracellular compartments^10,11,12^. After secretion, HMGB1 exerts inflammatory signals through activation of receptor for advanced glycation end-products (RAGE), toll-like receptor 2 (TLR2) or TLR4^10,13^. Since HMGB1-induced activation of TLRs has been observed in several disease conditions, or when challenged with LPS, the released HMGB1 facilitates delivery of LPS to TLR4, prompting endocytosis in the liver^14^. In contrast to this long-held notion, we show here that HMGB1 is highly expressed and retained in stromal cell nuclei in the pregnant uterus and is endowed with an anti-inflammatory role. Females with uterine deletion of *Hmgb1* show severe subfertility and give birth to small litters. One cause of this phenotype is perpetuating inflammatory conditions due to dysregulated migration of macrophages from the uterine stroma, leading to increased resorption rate and subfertility. These results unfold a novel function of HMGB1 in the pregnant uterus which has not been previously recognized.

## Results

### HMGB1 is expressed in the periimplantation uterus in a spatiotemporal manner

Hmg is a large family comprised of many members^15^. To identify major members in the uterus, we compared the expression levels of the family members by RNA-seq analysis of day 4 pregnant uteri and found that HMGB1 is one of the most abundant genes among the family members **(Fig. 1a)**. To examine the spatiotemporal expression of *Hmgb1* mRNA during the periimplantation period, we performed *in situ* hybridization using DIG method **(Fig. 1b, c)**. On day 1 through day 3, *Hmgb1* expression is predominantly localized in epithelial cells with some stromal localization on day 3. On the other hand, *Hmgb1* signals are primarily observed in stroma cells on days 4 and 5 (**Fig.1b, c**). Western blotting results show that HMGB1 protein levels decrease with pregnancy progression as assessed on days 4, 8 and 16 of pregnancy **(Fig. 1d)**. These results suggest that HMGB1 has specific functions in the uterus during early stages of pregnancy.

**Figure 1.**
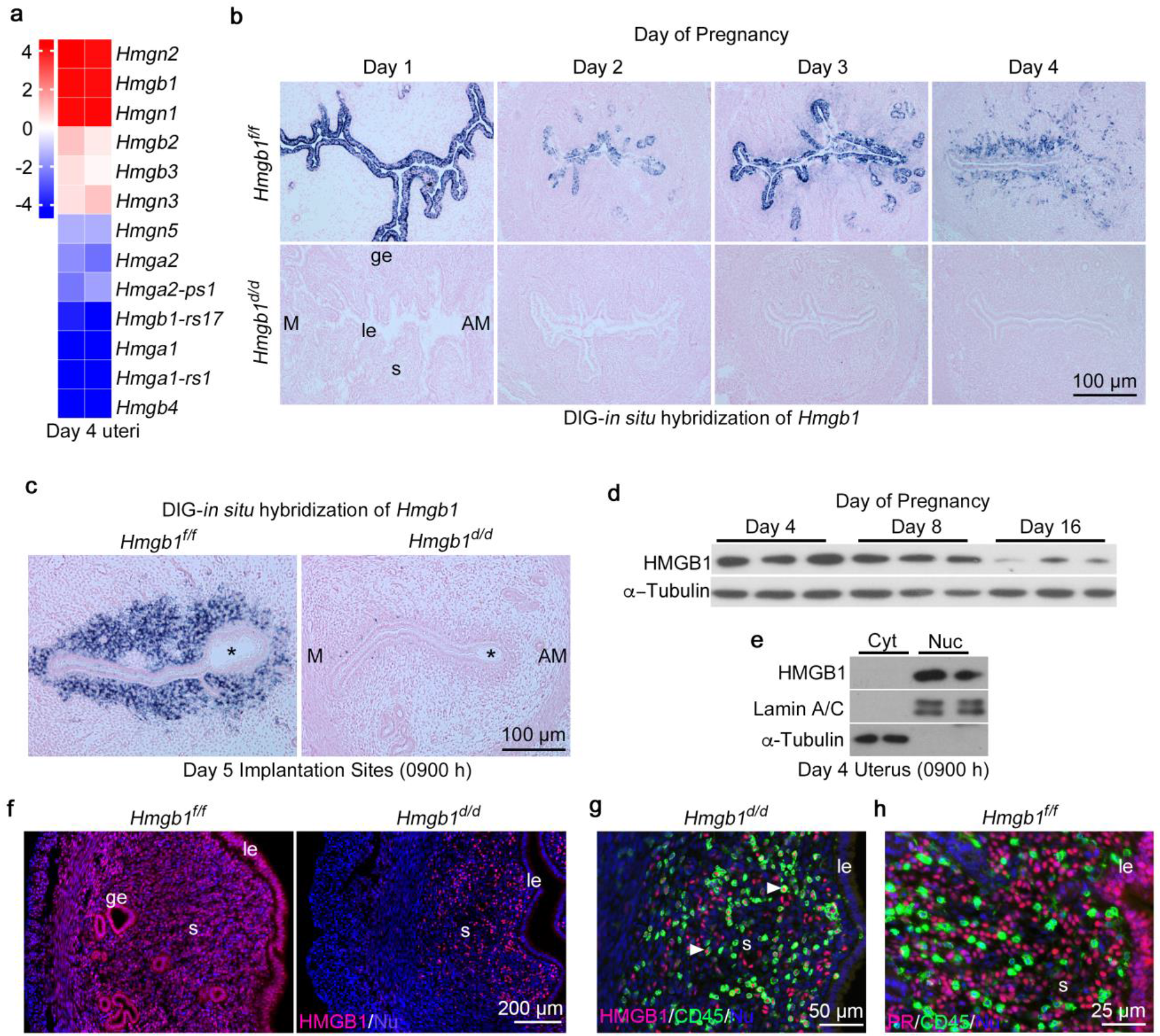
Expression of HMGB1 in the Early Pregnant Uteri. **a**, Heatmap shows relative expression levels of *Hmg* family members in day 4 pregnant uteri. Data are expressed as reads per kilobase per million (RPKM) for RNA-seq analysis, n=2. **b, c**, *In situ* hybridization of *Hmgb1* expression in *Hmgb1*^*f/f*^ and *Hmgb1*^*d/d*^ uteri from days 1-5 of pregnancy. Asterisks indicate blastocysts. le: luminal epithelium, s: stroma, ge: glandular epithelium. M: mesometrial pole, AM: antimesometrial pole. Scale bar: 100 μm. **d**, Western blotting of HMGB1 in uterine tissues on days 4, 8 and 16 of pregnancy. α-Tubulin was used as a loading control. **e**, Hmgb1 levels in the cytosolic and nuclear fractions from day 4 pregnant uteri. Lamin A/C and α-Tubulin are markers for nuclei and cytosol, respectively. Cyt: Cytosol, Nuc: Nuclei. **f**, IF of HMGB1 in day 4 uteri of *Hmgb1*^*f/f*^ and *Hmgb1*^*d/d*^ mice. le: luminal epithelium, s: stroma, ge: glandular epithelium. Scale bar: 200 μm. **g**, IF of HMGB1 and CD45 in day 4 uteri of *Hmgb1*^*d/d*^ mice. le: luminal epithelium, s: stroma. Asterisks indicate colocalization of CD45 and HMGB1. le: luminal epithelium, s: stroma. Scale bar: 50 μm. **h**, IF of PR and CD45 in day 4 uteri of *Hmgb1*^*f/f*^ mice. le: luminal epithelium, s: stroma, ge: glandular epithelium. le: luminal epithelium, s: stroma. Scale bar: 25 μm. Each image is a representative from at least 3 independent experiments.

HMGB1 protein is primarily localized in the nucleus as evident from Western blotting results of isolated nuclei and immunostaining **(Fig. 1e, f)**. The broader expression of HMGB1 protein than mRNA in floxed uteri could be due to differential stability of mRNA versus protein **(Fig. 1b, f)**. In vitro studies showed that HMGB1 is released from nuclei under certain stress conditions in the lung and liver^13^. However, we do not see translocation of uterine HMGB1 from the nucleus to the cytoplasm in mice even after LPS stimulation **(Supplementary Figure 1a)**. These observations suggest that uterine HMGB1 executes its functions engaging nuclear activity under in vivo settings.

### Mice with uterine deletion of *Hmgb1^d/d^* show subfertility due to aberrant implantation

To explore the function of HMGB1 in pregnant uteri, we generated mice with uterine deletion of *Hmgb1* (*Hmgb1^d/d^*) by crossing *Hmgb1* floxed mice (*Hmgb1^f/f^*) with progesterone receptor-Cre transgenic mice (*Pgr^Cre/+^*)^16,17^. These mice show efficient deletion of *Hmgb1* in the pregnant uterus at both mRNA and protein levels **(Fig. 1b, c, f and Supplementary Figure 1b, c)**. In deleted uteri on day 4, many scattered cells are still HMGB1-positive in the stromal bed on day 4 **(Fig. 1f)**. Co-staining of HMGB1 and CD45 (marker of leukocytes) in the deleted uteri shows that scattered HMGB1-positive cells are CD45 positive, providing evidence that HMGB1 is also expressed in leukocytes and are not deleted by *Pgr-Cre* (**Fig. 1g**). To show if these CD45 positive leucocytes are PR negative, PR and CD45 were co-stained in day 4 *Hmgb1*^*f/f*^ uteri (**Fig. 1h**). These results implicate that PR is primarily expressed in uterine stromal cells, but not leucocytes, while HMGB1 is expressed in both stromal cells and leucocytes.

The above results provoked us to explore the pregnancy outcome in this uterine HMGB1 deleted (*Hmgb1^d/d^*) females. We found that deleted females show severe subfertility: only 39% of plug-positive *Hmgb1*^*d/d*^ females produce litters and the litter size is significantly smaller than those in littermate floxed mice **(Fig. 2a, b)**. No differences are noted in birth timing between the two genotypes **(Fig. 2c)**. Interestingly, about 50% of pups born from *Hmgb1*^*d/d*^ females died within 2 days after birth **(Fig. 2d)**, suggesting unfavorable conditions in the *Hmgb1*^*d/d*^ pregnant womb.

**Figure 2.**
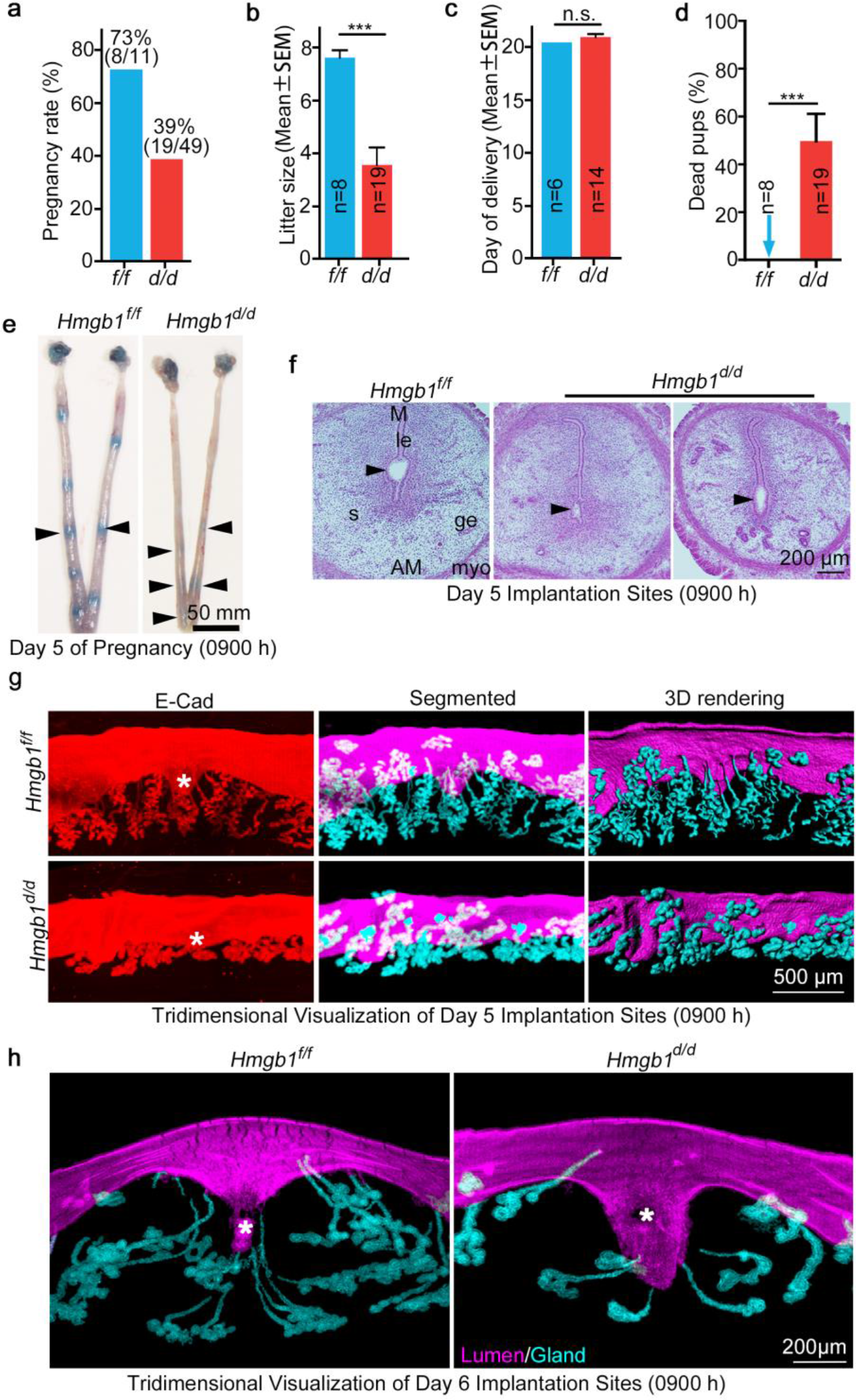
Deletion of *Hmgb1* in Uteri Compromises Embryo Implantation and Pregnancy Outcome. **a**, Percentage of pregnancy rate in *Hmgb1*^*f/f*^ and *Hmgb1*^*d/d*^ females. The number within brackets indicate females with pups over total number of plug-positive females. **b**, Average number of litter sizes in *Hmgb1*^*f/f*^ and *Hmgb1*^*d/d*^ females. *Hmgb1*^*f/f*^ (n= 8); *Hmgb1*^*d/d*^ (n= 19). Data are presented as mean ± SEM, \*\*\**P* < 0.001 (student’s *t*-test). **c**, Average day of delivery. n = 6 for *Hmgb1*^*f/f*^ and n = 14 for *Hmgb1^d/d^*. Data are presented as mean ± SEM, n.s.: not significant (student’s *t*-test). **d**, Percentage of dead pups in *Hmgb1*^*f/f*^ and *Hmgb1*^*d/d*^ females. \*\*\**P*< 0.001. **e**, Day 5 implantation sites (blue bands) in *Hmgb1*^*f/f*^ and *Hmgb1*^*d/d*^ females. Arrowheads indicate weak blue bands. Scale bar: 50 mm. **f**, Histology of day 5 implantation sites in *Hmgb1*^*f/f*^ and *Hmgb1*^*d/d*^ females. Arrowheads indicate the location of blastocysts. le: luminal epithelium, s: stroma, ge: glandular epithelium, myo: myometrium. M: mesometrial pole, AM: antimesometrial pole. Scale bar: 200 μm. **g**, 3D imaging of day 5 implantation sites from *Hmgb1*^*f/f*^ and *Hmgb1*^*d/d*^ females. Staining of E-Cad (epithelial cell marker), segmented glands and uterine lumen and 3D rendering of day 5 implantation site. Images were generated by a Nikon A1R Multiphoton Microscope with LWD 16X water objective with 3 μm Z-stack. Asterisks indicate embryos. Scale bar: 500 μm. **h**, 3D visualization of day 6 implantation sites from *Hmgb1*^*f/f*^ and *Hmgb1*^*d/d*^ females. IF of E-Cad was performed to visualize LE and glands. Images were generated as described in **g**. Asterisks indicate embryos. Scale bar: 200 μm. Each image is a representative from at least 3 independent experiments.

To compare stage-specific pregnancy phenotypes, mice were intravenously injected with blue dye solution on day 5. We found that *Hmgb1*^*d/d*^ uteri have faint blue bands as compared to littermate floxed uteri **(Fig. 2e)**. Histological analysis shows inferior implantation chambers in *Hmgb1*^*d/d*^ mice **(Fig. 2f)**. To determine the crypt (implantation chamber) formation and gland-crypt assembly for direct communication with the implanting blastocyst within the chamber, tridimensional (3D) visualization was employed after fixing and tissue clearing^6^. Days 5 and 6 floxed uteri showed well-defined spear-shaped implantation crypts with drawn-out and well-developed glands. These phenotypes are not observed in most *Hmgb1*^*d/d*^ mice with poor implantation **(Fig. 2g, h)**.

The aberrant implantation on days 5 and 6 led us to assess the uterine receptivity in these mice. Expression levels of uterine receptivity markers *Ihh* (P_4_ responsive) and *Msx1* (hormone unresponsive) genes^1,3^ are comparable between *Hmgb1*^*f/f*^ and *Hmgb1*^*d/d*^ mice **(Fig. 3a)**. In contrast, P_4_-dependent *Hoxa10*^18^, which is expressed in stromal cells and is thought to prepare stromal cell to decidualization is down-regulated in *Hmgb1*^*d/d*^ uteri (**Fig. 3a**). These results suggest that epithelial-stromal cross-talk in uterine receptivity is compromised by *Hmgb1*-deletion. We also examined if serum levels of P_4_ and expression patterns of progesterone receptor (PR) and estrogen receptor (ERα) are altered between *Hmgb1*^*f/f*^ and *Hmgb1*^*d/d*^ uteri, and found that these parameters are comparable between the two groups **(Supplementary Figure 2a-c)**. Blood vessel distribution as assessed by CD31 staining appears unaltered **(Supplementary Figure 2c)**. The expression of FOXA2, a critical transcription factor for uterine gland formation and function^6,19^, is also not affected in *Hmgb1*^*d/d*^ uteri **(Supplementary Figure 2d)**.

**Figure 3.**
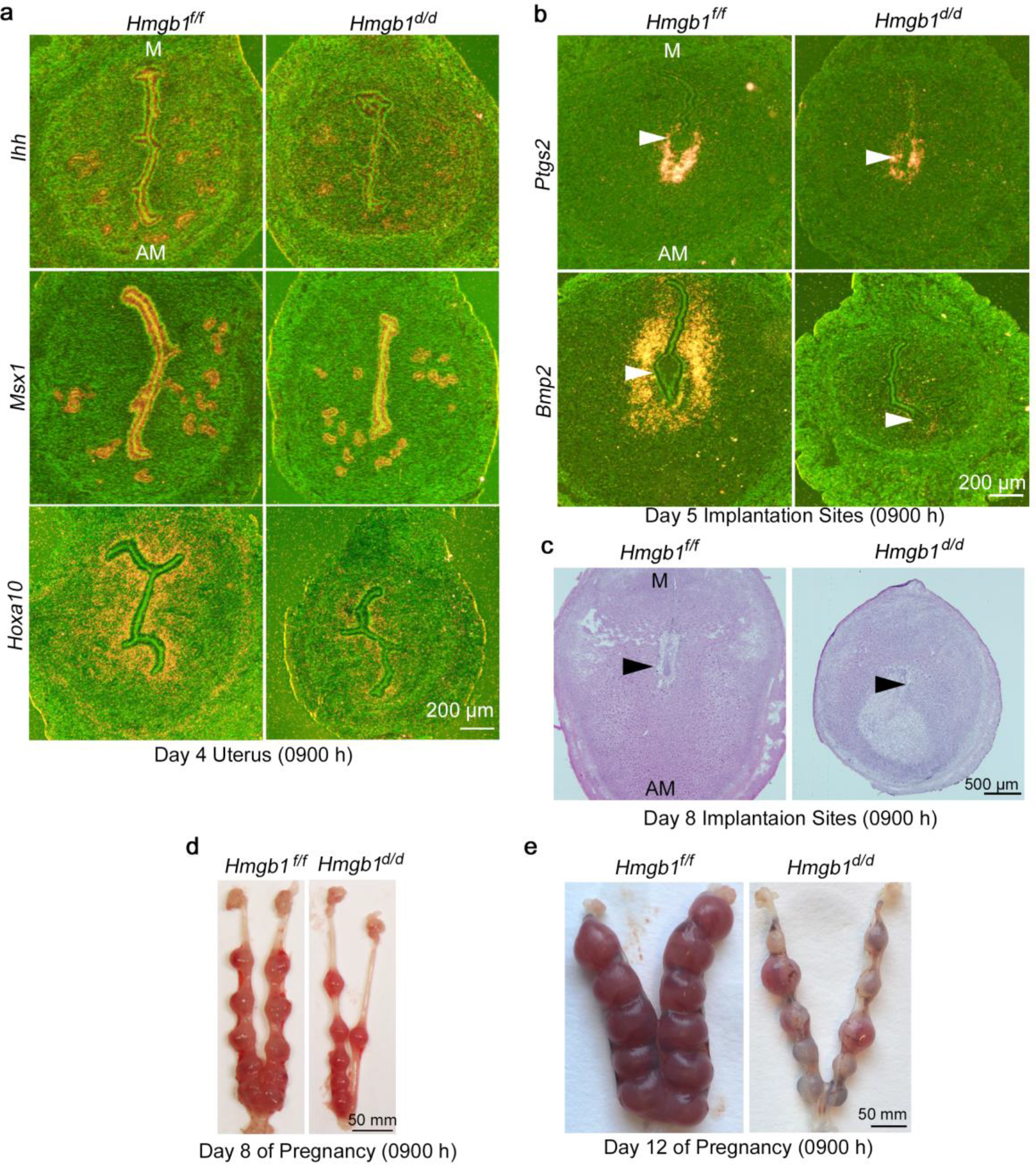
HMGB1 Deletion in Uteri Causes Abnormal Decidualization. **a,** *In situ* hybridization of *Msx1*, *Ihh* and *Hoxa10* in day 4 pregnant uteri from *Hmgb1*^*f/f*^ and *Hmgb1*^*d/d*^ females. M: mesometrial pole, AM: antimesometrial pole. Scale bar: 200 μm. **b**, *In situ* hybridization of *Ptgs2* and *Bmp2* in day 5 implantation sites from *Hmgb1*^*f/f*^ and *Hmgb1*^*d/d*^ females. Arrowheads indicate blastocysts. M: mesometrial pole, AM: antimesometrial pole. Scale bar: 200 μm. **c**, Histology of day 8 implantation sites from *Hmgb1*^*f/f*^ and *Hmgb1*^*d/d*^ females. Arrowheads indicate embryos. M: mesometrial pole, AM: antimesometrial pole. Scale bar: 500 μm. **d**, Day 8 pregnant uteri from *Hmgb1*^*f/f*^ and *Hmgb1*^*d/d*^ females. Scale bar: 50mm. **e**, Day 12 pregnant uteri from *Hmgb1*^*f/f*^ and *Hmgb1*^*d/d*^ females. Scale bar: 50 mm. Each image is a representative from at least 3 independent experiments.

The attachment reaction seems to be defective as indicated by diminished *Ptgs2* (encoding cyclooxygenase 2, Cox2) expression with profound decrease in the expression of bone morphogenetic protein (*Bmp2*), a marker for decidualization, in day 5 *Hmgb1*^*d/d*^ uteri **(Fig. 3b)**^1,20,21^. Consistent with these findings, the implantation sites on day 8 are much smaller in *Hmgb1*^*d/d*^ mice **(Fig. 3c, d)**. These smaller implantation sites eventually result in resorptions on day 12 (**Fig. 3e**). Taken together, these results show that uterine deletion of *Hmgb1* compromises embryo implantation and pregnancy success.

### *Hmgb1*^d/d^ females show increased uterine accumulation of macrophages

Since HMGB1 is thought to induce inflammatory signals^10,13^, we examined immune cell distribution in uteri of *Hmgb1*^d/d^ mice and found higher enrichment of leukocytes (CD45-positive cells) in the uterine stroma lacking *Hmgb1* on day 4 of pregnancy **(Fig. 4a)**. This analysis was followed by visualization of macrophage population in the stroma by IF of F4/80, a widely-used macrophage marker. Compared to *floxed* uteri, *Hmgb1*^*d/d*^ uteri show significant increases in macrophage population on days 4 and 5 of pregnancy (**Fig. 4b** **and** **Supplementary Figure 3a**). Tridimensional visualization of F4/80 and E-Cadherin (E-Cad) co-staining with sectional view of day 5 uteri further reinforced that more macrophages are enriched in the subluminal region in *Hmgb1*^*d/d*^ mice (**Fig. 4c** **and Videos S1, S2).** We noted that some macrophages invaded through epithelial cells and adhered to embryos **(Fig. 4c, d)**. Consistent with the staining of CD45 and PR, these macrophage in day 4 *Hmgb1*^*d/d*^ uterus is also PR negative **(Fig. 4e)** which indicates that the increased enrichment of macrophages is primarily attributed to stromal *Hmgb1* depletion, but not due to its deficiency in leucocytes. Sparse distribution of neutrophils and NK cells in day 5 pregnant uteri supports that increased accumulation of macrophages primarily drives the hostile environment to embryo implantation **(Supplementary Figure 3b, c)**. This is a potential reason for reduced number of implantation sites and litter sizes **(Fig. 2b, e)**. Notably, *Hmgb1* floxed and deleted mice show comparable number of leukocytes and macrophages in the ovary **(Supplementary Figure 3d)**.

**Figure 4.**
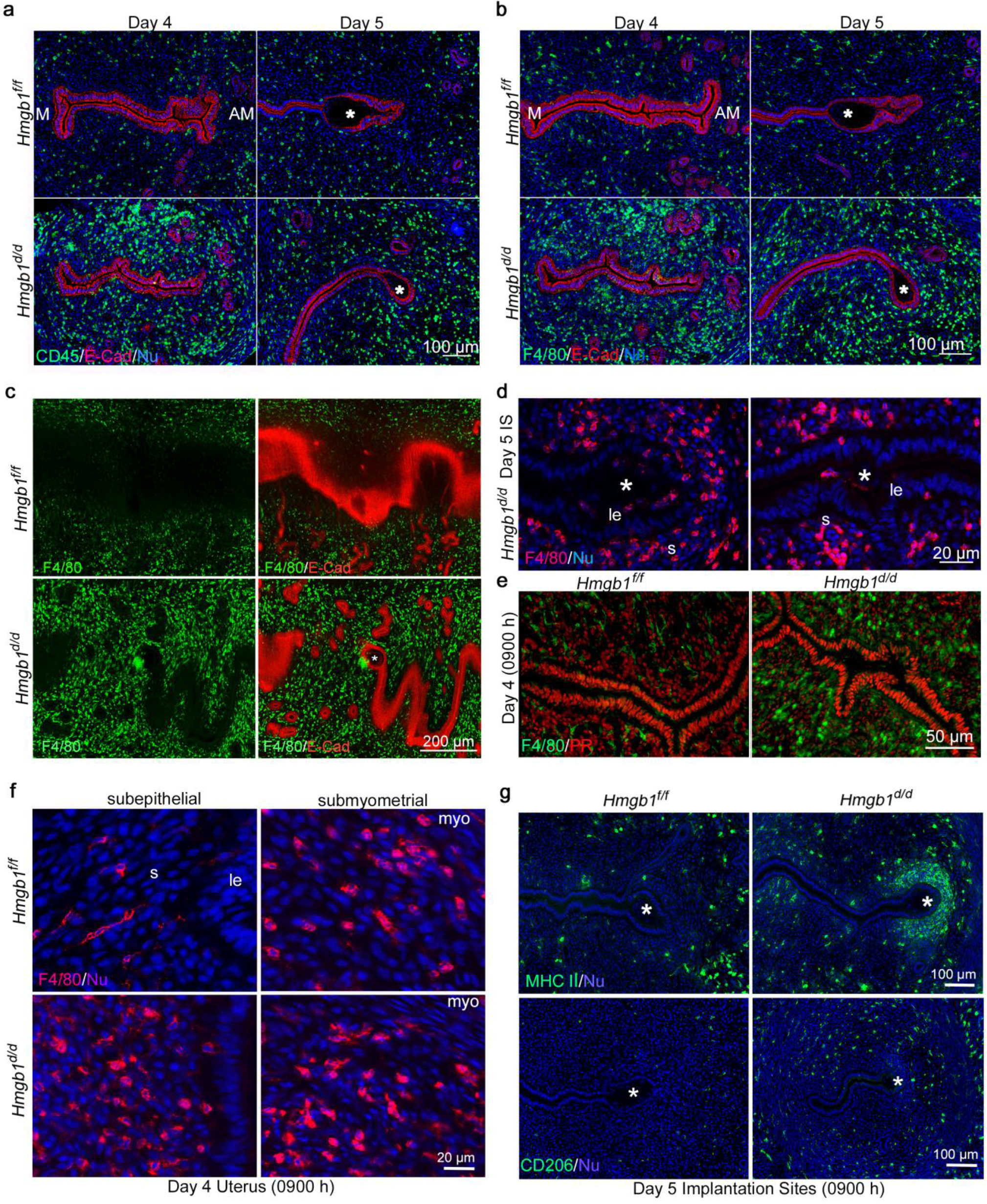
Increased M1-Macrophages in *Hmgb1*^*d/d*^ Pregnant Uteri. **a,** IF of CD45 (pan-leukocytes marker) and E-Cad in days 4 and 5 pregnant uteri from *Hmgb1*^*f/f*^ and *Hmgb1*^*d/d*^ females. Asterisks indicate blastocysts. M: mesometrial pole, AM: antimesometrial pole. Scale bar: 100 μm. **b**, IF of F4/80 (macrophage marker) and E-Cad in days 4 and 5 pregnant uteri of *Hmgb1*^*f/f*^ and *Hmgb1*^*d/d*^ females. Asterisks indicate blastocysts. M: mesometrial pole, AM: antimesometrial pole. Scale bar: 100 μm. **c**, Sectional view of 3D visualization of E-Cad and F4/80 staining in day 5 uteri in *Hmgb1*^*f/f*^ and *Hmgb1*^*d/d*^ females. Asterisks indicate blastocysts. Scale bar: 200 μm. **d,** IF of F4/80 in day 5 pregnant uteri from *Hmgb1*^*d/d*^ females. Invasion of macrophages into the implantation chamber (crypt) was observed. Asterisks indicate blastocysts. le, luminal epithelium. s, stroma. Scale bar: 20 μm. **e,** IF of F4/80 and PR in day 4 pregnant uteri from *Hmgb1*^*f/f*^ and *Hmgb1*^*d/d*^ females. Scale bar: 50 μm. **f**, IF of F4/80 in day 4 pregnant uteri from *Hmgb1*^*f/f*^ and *Hmgb1*^*d/d*^ females. Subepithelial and submyometrial regions are shown. le: luminal epithelium, s: stroma, myo: myometrium. Scale bar: 20 μm. **g**, IF of MHC II (M1 macrophage marker) and CD206 (M2 macrophage marker) in day 5 *Hmgb1*^*f/f*^ and *Hmgb1*^*d/d*^ pregnant uteri. Asterisks indicate blastocysts. Scale bar: 100 μm. Each image is a representative of at least 3 independent experiments.

### Macrophage polarization is dysregulated in *Hmgb1^d/d^* uteri

Histological analysis show that the shape of macrophages is different depending on their location: elongated macrophages are located in the subluminal stroma compared to oval-shaped (roundish) macrophages in the submyometrial stroma in *floxed* day 4 uteri **(Fig. 4e)**. In contrast, roundish macrophages are present throughout the endometrium in *Hmgb1*^*d/d*^ females **(Fig. 4e)**. These roundish macrophages are also seen in the stroma close to the implantation site of *Hmgb1*^*d/d*^ on day 5, whereas macrophages are virtually absent at the implantation site of *Hmgb1*^*f/f*^ mice **(Fig. 4b, c)**.

The differences in macrophage shape prompted us to ask if there are different subtypes of macrophage populations and their functional differences in the uterus. Macrophages are classified primarily into two subtypes, pro-inflammatory (M1: MHC II positive) and anti-inflammatory (M2: CD206 positive) types^22,23,24^. Since the shape of macrophages influences their phenotypes^24^, we evaluated cell types of M1 and M2 macrophages. In *Hmgb1*^*f/f*^ uteri on day 5, elongated M1-type and roundish M2-type macrophages are located at the submyometrial region with no macrophages around the implanting blastocyst **(Fig. 4f)**. In contrast, *Hmgb1*^*d/d*^ uteri showed significant accumulation of roundish M1 macrophages in the vicinity of blastocyst like a “necklace”. M2 macrophage distribution is very sparse in day 5 floxed uteri, whereas the number was still higher in the deleted mice. This may be due to restrain pro-inflammatory responses arising from increased numbers of M1 macrophages **(Fig. 4f)**.

We found uterine HMGB1 suppresses pro-inflammatory condition during periimplantation period. This is in contrary to findings of secreted HMGB1^10,13,15^. This raises an interesting question as to how nuclear HMGB1 mediates macrophage migration in pregnant uteri.

### HMGB1-mediated PR activation is critical for uterine receptivity

Since the establishment of early pregnancy is accompanied with dynamic interplay of maternal hormones, the defective implantation and abnormal macrophage enrichment in HMGB1 ablated mice led us to explore the effects of hormones on macrophages distribution from days 1 to 4 due to the remarkable switch of the uterine milieu from estrogen to progesterone dominance. We observed that macrophages progressively migrate away from the stroma from day 3 onwards in the floxed uterus, retaining a relatively small number on day 4 **(Fig. 5a)**. In contrast, *Hmgb1* deficient uteri on days 3 and 4 consistently show elevated number of macrophages **(Fig. 5a)**. These results also suggest that HMGB1 mediated timely display of macrophage distribution under the influence of the maternal hormone is critical for successful implantation. There is evidence that monocytes, precursors of macrophages, are enriched and propagated in the myometrium^25^. To exclude the possibility that the increased population of macrophages in *Hmgb1*^*d/d*^ uteri is the result of proliferation of resident macrophages, we performed co-immunostaining of F4/80 and Ki67. Failure to observe their co-localization supports that increased accumulation of macrophages is primarily due to either their increased trafficking in the stroma or delayed migration from the stroma, but not proliferation **(Supplementary Figure 3e)**. These results suggest different mechanisms for maintaining tissue-specific macrophage densities. The increased macrophage accumulation was also observed in *Hmgb1*^*d/d*^ uteri on day 4 of pseudopregnancy **(Supplementary Figure 3f)**, suggesting that this abnormality is caused by an aberration of uterine milieu, but not due to the presence of blastocysts in the uterus.

**Figure 5.**
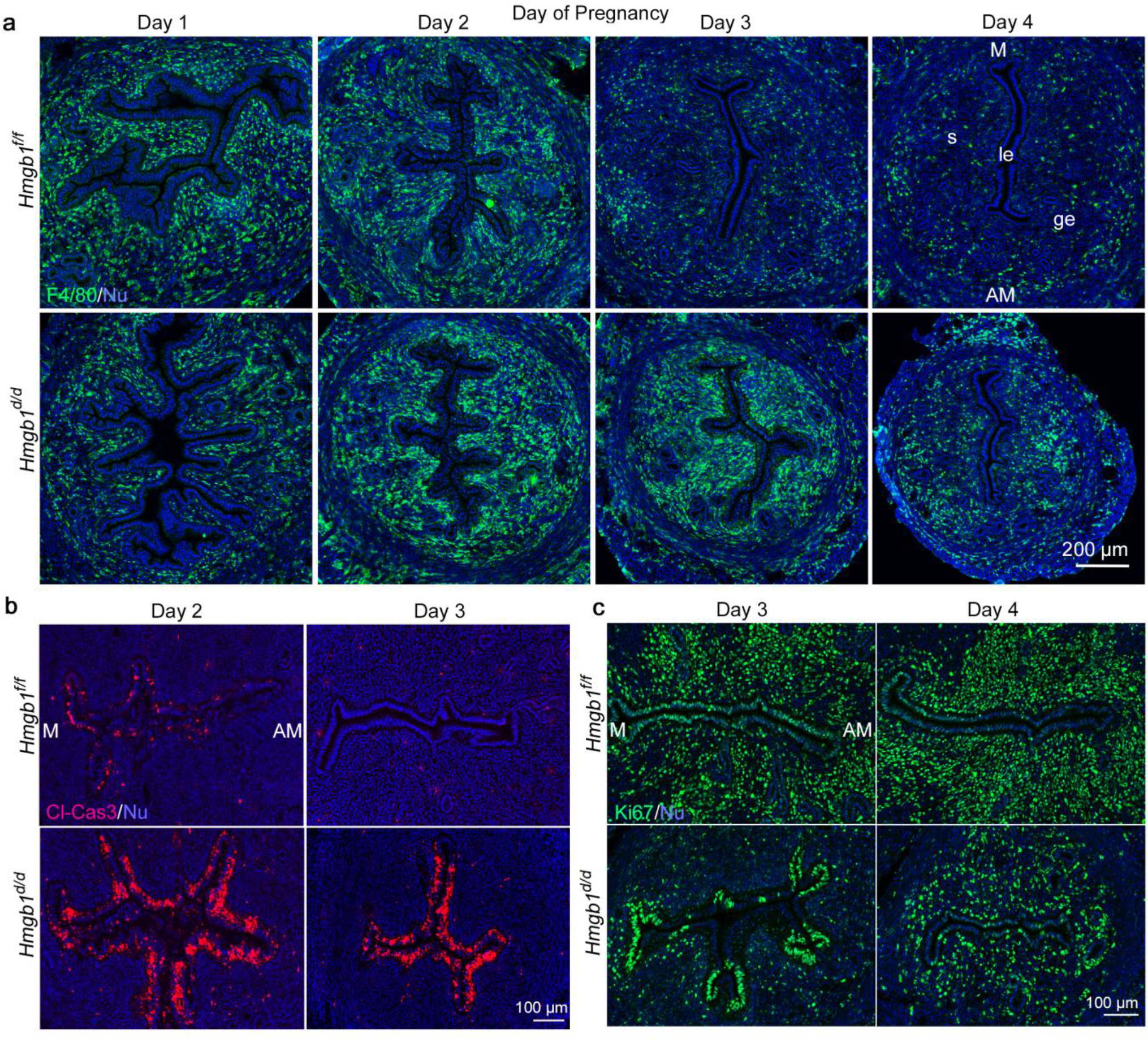
Deletion of *Hmgb1* in Uteri Results in Abnormal Macrophage Infiltration and Stromal Cell Proliferation. **a**, IF of F4/80 in days 1-4 pregnant uteri from *Hmgb1*^*f/f*^ and *Hmgb1*^*d/d*^ females. le: luminal epithelium, s: stroma, ge: glandular epithelium. M: mesometrial pole, AM: antimesometrial pole. Scale bar: 200 μm. **b**, IF of cleaved Caspase-3 (apoptotic cell marker) in days 2 and 3 pregnant uteri from *Hmgb1*^*f/f*^ and *Hmgb1*^*d/d*^ females. Casp3: Cleaved Caspase-3, M: mesometrial pole, AM: antimesometrial pole. Scale bar: 100 μm. **c**, IF of Ki67 in days 3 and 4 pregnant uteri from *Hmgb1*^*f/f*^ and *Hmgb1*^*d/d*^ females. M: mesometrial pole, AM: antimesometrial pole. Scale bar: 100 μm. Each image is a representative from at least 3 independent experiments.

Changing levels of ovarian hormones estrogen and progesterone direct the establishment of uterine receptivity by guiding cell type specific proliferation and differentiation during days 1-4 of pregnancy^1,2^. Compared with day 2, the day 3 uterus is marked by the termination of apoptosis in epithelial cells with the onset of stromal cell proliferation under the influence of increasing P_4_ levels from the newly formed corpora lutea^2,3,26^. Increased macrophage accumulation in *Hmgb1*^*d/d*^ uteri on day 3 **(Fig. 5a)**, led us to presume that transition of uterine cells between days 2 and 3 is compromised in deleted uteri. The evidence that *Hmgb1*^d/d^ uteri on day 3 show sustained apoptosis in epithelial cells as indicated by distinct staining of cleaved Caspase-3 supports this assumption **(Fig. 5b)**. This abnormal cell death might be one cause of increased immune cells in *Hmgb1*^*d/d*^ uteri as indicated by the previous observation that *Hmgb1* deficiency causes cell damage and death, resulting in tissue inflammation ^27^. This aberrant apoptosis is followed by reduced stromal cell proliferation as evident from reduced number of Ki67-positive stromal cells on days 3 and 4 in *Hmgb1*^*d/d*^ females **(Fig. 5c)**. Collectively, our results show that an abrogated transition of the uterus from day 2 to 3 in *Hmgb1*^d/d^ females disturbs uterine receptivity and thus embryo implantation.

Since poor P_4_ responsiveness accounts for abnormal cell proliferation^1,26^, we asked if PR activation is affected in *Hmgb1*-deleted uteri. In the uterus, PR has been reported to be critical for anti-inflammation: activation of PR by its specific agonist inhibits macrophage invasion into the endometrium^28^. In addition, PR knockout females are completely infertile^29^. HMGB1 can bind to PR and enhance their activation^30,31^. To assess PR activation in the presence or absence of HMGB1, we performed luciferase reporter assays using primary stromal cells from floxed and deleted mice transfected with PRE2-Tk-Luc. We found that PR activation is substantially reduced in *Hmgb*^*d/d*^ stromal cells compared to both of *Hmgb1*^*f/f*^ and *Pgr*^*Cre/+*^ cells **(Fig. 6a)**. In contrast, activation of glucocorticoid receptor, which is also reported to interact with HMGB1^31,32^, is comparable between both genotypes **(Supplementary Figure 4a)**, indicating that HMGB1 interacts with PR in the uterus. To determine if insufficient PR signaling in *Hmgb1*^d/d^ mice contributes to implantation failure, exogenous P_4_ (2 mg/mouse) was injected to *Hmgb1*^*d/d*^ females as we have previously reported^33^. We found this treatment failed to rescue defective implantation **(Fig. 6b)**. In addition, accumulation of M1 macrophages is not neutralized by P_4_ treatment to *Hmgb1*^*d/d*^ females **(Supplementary Figure 4b)**. These results suggest that HMGB1’s role in PR signaling pathway is more complex and cannot be substituted by P_4_ injection.

**Figure 6.**
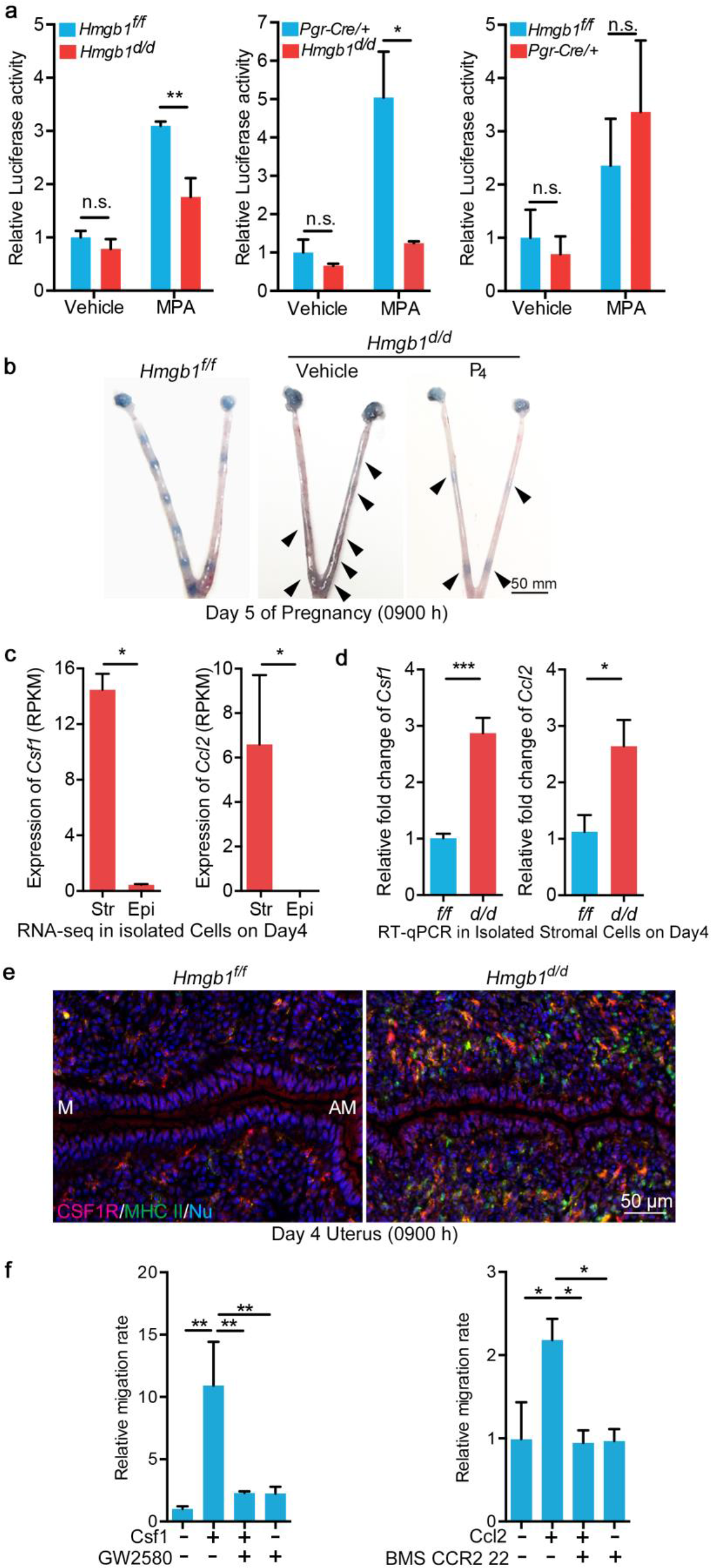
*Hmgb1* Deleted Stromal Cells Cause Dysregulated Inflammatory Signals. **a**, Decreased PR activation in *Hmgb1*^*d/d*^ stromal cells determined by PRE2-Tk-luciferase assay. n = 2-4 for each genotype. Data are presented as mean ± SEM, \**P* < 0.05, \*\**P* < 0.01 and n.s.: not significant by one-way ANOVA. **b**, Day 5 implantation sites (blue bands) in P_4_-supplemented *Hmgb1*^*d/d*^ females. Arrowheads indicate weak blue bands. Scale bar: 50 mm. **c**, Expression levels of macrophage attractants, *Csf1* and *Ccl2*, were analyzed by RNA-seq analysis in isolated epithelial/stromal cells from day 4 uteri. n = 3 for each group. Str: Stroma, Epi: Epithelium. Data are presented as mean ± SEM, \**P* < 0.05 and \*\*\**P* < 0.005 by Student *t*-test. **d**, Quantitative RT-PCR showed increased levels of macrophage attractants (*Ccl2* and *Csf1*). n = 4 for each genotype. Data are presented as mean ± SEM, \**P* < 0.05 and \*\*\**P* < 0.005 by student’s *t*-test. **e**, IF of CSF1R and MHC II in day 4 pregnant uteri from *Hmgb1*^*f/f*^ and *Hmgb1*^*d/d*^ females. M: mesometrial pole, AM: antimesometrial pole. Scale bar: 50 μm. Each image is a representative from at least 3 independent experiments. **f**, Migration assays show that Csf1 and Ccl2 attract macrophages depending on their specific receptors. GW2580 and BMS CCR2 22 were used as Csf1 and Ccl2 receptor antagonists, respectively. These assays were performed in three wells for each group and three fields/wells were quantified. Data are presented as mean ± SEM, \**P* < 0.05 and \*\**P* < 0.01 (One-way ANOVA).

### Macrophage attractants are upregulated in *Hmgb1* deleted stroma

The above results led us to ask the mechanism by which macrophages accumulate and are retained in the stroma. Macrophages migration is regulated by chemokines; Ccl2 and Csf1 are considered important macrophage attractants^22^. As described above, *Hmgb1*^*d/d*^ uteri have reduced expression of *Hoxa10* (**Fig. 3a**), a P_4_-responsive stromal gene, and is critical for stromal cell proliferation^18^. In this context, deletion of *Hox* genes including *Hoxa10* causes increased expressions of cytokines, promoting macrophage accumulation in the mouse uterus^34^. These findings prompted us to examine the level of chemokines in *Hmgb1* deleted uteri. Our RNA-Seq analysis in isolated epithelial and stromal cells from day 4 uteri show that the expression of these attractants are higher in stromal cells than epithelial cells (**Fig. 6c**). This is in contrast to previous report of *Csf1 mRNA* localization *in situ* hybridization in uterine epithelial cells on day 4^35^. This disparity between these two studies could be due to genetic background of mice and/or environmental and dietary composition^33,36^. The data from single cell RNA-Seq analysis of the uterus^37^ also reveals that *Ccl2* and *Csf1* are highly expressed in specifically stromal cells with exclusive expression of *Csf1r* in macrophages (**Supplementary Figure 4c**). To obtain further insight, we performed qRT-PCR to assess cytokine levels in isolated purified stromal cells after removing macrophages. We observed that *Csf1* and *Ccl2* are significantly upregulated in *Hmgb1*^*d/d*^ stromal cells **(Fig. 6d)**. These results corroborate with a higher level of CSF1R-positive M1 macrophages in *Hmgb1*-deleted uteri **(Fig. 6e)**. With these results in hand, we examined if these chemokines serve as attractants using a macrophage cell line *in vitro* assays. Indeed, CSF1 was more potent than CCL2 to attract macrophages **(Fig. 6f).** These observations mechanistically indicate that *Hmgb1*-deleted uteri sustain macrophages in the stroma due to higher levels of attractants and reduced PR activation. Furthermore, our results suggest that HMGB1 regulates macrophage trafficking through CSF1 and CCL2 in the uterus conducive to blastocyst implantation.

## Discussion

The highlight of this investigation is that nuclear HMGB1 in the uterus is essential to pregnancy success in mice. HMGB1 limits macrophage accumulation in the uterus by maintaining PR activation, restraining chemokine levels, and encouraging retention of uterine macrophages in the stroma. Our present observations are in stark contrast to the dogma that HMGB1 serves as an inflammatory ligand in many organs. The unique and programmed mixture of stromal cells and leukocytes in the uterus fine-tunes the pregnancy progress by orchestrating inflammation homeostasis. After mating, a large number of leukocytes including macrophages accumulate in the uterus influenced by estrogen and seminal fluid, evoking inflammatory-like responses^38^. This accumulation of immune cells begins to migrate away from the stroma from day 3 onward and gradually assembles in the myometrium^8,39^. It is thought that this gradual migration from the stroma is critical for successful pregnancy^39,40^. Although the importance of deciduae and placentas for immune suppression has repeatedly been described^40,41^, it remains elusive how immune cell infiltration in the uterus is subdued prior to and during implantation.

Previous results of in vitro studies and recently generated conditional knockout mice show the complexities of HMGB1’s functions that are context and tissue dependent^16,42^. Our in vivo studies showing that uterine-specific deletion of *Hmgb1* leads to the demise of blastocyst implantation or defective implantation, creating adverse ripple effects throughout the course of pregnancy is a previously undocumented role of HMGB1. Our results also show that uterine distribution and polarization of macrophages are under the influence of HMGB1. The observation of elongated macrophages (M1) in the subepithelial region and roundish macrophages (M2) in the submyometrial stroma in floxed uteri suggest that the uterine microenvironment regulates polarization of macrophages from M1 to M2 stage. Skewed distribution of M1 and M2 macrophages toward M1 macrophages in the stroma surrounding the implanting blastocyst in the absence of HMGB1 provides evidence that HMGB1 is critical to the polarization and distribution of macrophages conducive to implantation. Taken together, these observations suggest that HMGB1 has critical roles to regulate macrophage migration and polarization in the pregnant uteri.

Perhaps the dysregulated uterine microenvironment is a cause of this aberrant macrophage accumulation in the absence of HMGB1. In mice, apoptosis of epithelial cells is prominent on day 2 but subsides on day 3 with initiation of stromal cell proliferation and migration of macrophages toward the myometrium^1,2,3^. The sustained epithelial cell death that continues in *Hmgb1*^*d/d*^ uteri perhaps alters the uterine environment toward inflammation that leads to reduced stromal proliferation.

HMGB1 influences the levels of inflammatory cytokines in stromal tissues. The purified stromal cells from *Hmgb1*^*d/d*^ uteri indeed show higher expression levels of macrophage attractants. These results suggest that immune cells, primarily macrophages, are hyper-activated in the endometrium deficient in HMGB1. Collectively, these observations implicate that uteri lacking HMGB1 sustain inflammatory milieu during the early pregnancy. Recently a group reported *Hmgb1* expression in the uterus by in situ hybridization without any in vivo genetic evidence of its function in pregnancy; this group claims that HMGB1 plays a role in decidualization as assessed *in vitro* studies^43^. In contrast, our study presents genetic evidence that HMGB1 is essential for early pregnancy events prior to and during blastocyst implantation. We believe that defective decidualization is a consequence of the derailed implantation process, creating adverse ripple effects^5,44,45^. Although HMGB1 could be important for decidualization, decidua-specific deletion of *Hmgb1* is required to address this issue which is not achievable at present due to lack of decidua-specific Cre driver.

It is also striking finding that uterine HMGB1 is working mainly in the nuclei but not as a secreted molecule. Many studies focused on HMGB1’s function as a secreted ligand for TLRs or RAGE^10,12,13,15,46,47^. However, our study shows that HMGB1 is stable in the nucleus, even after exposure to LPS. HMGB1’s interaction with steroid hormone receptors including PR and GR supports the gene regulatory role of nuclear HMGB1^30,31,48^. In fact, Dean and his associates have shown that HMGB1 increases the DNA-binding affinity of PR and other steroid receptors. Carboxyl terminal extension of PR contains a specific binding site for HMGB that is critical for stimulating PR-PRE binding activity^49^. Our observations of reduced PR activity but not GR reveal that the interaction between HMGB1 and nuclear receptor is tissue specific and context dependent. The landscape of HMGB1 function in whole genome is yet to be determined^50^.

P_4_ injection was used to rescue embryo implantation and decidualization attributed by reduced PR signaling^33,51^, but failure of P_4_ administration to rescue embryo implantation in *Hmgb1*^d/d^ mice indicates that the role HMGB1 is not replaceable by P_4_ supplementation in the presence of depressed PR activation. This is compatible with the finding that serum levels of P_4_ are comparable between floxed and deleted mice on day 4. Whether HMGB1 is a co-regulator of PR in uterus requires further investigation.

Macrophages are retained in day 3 uteri even with the comparable P_4_ levels in both WT and ablated mice. This result places the significance of HMGB1 as a critical regulator of macrophage migration. Csf1 and Ccl2 are considered very important attractants for macrophage migration^22^. Deficient Csf1 levels in *Csf1*^*op/op*^ mice was shown to reduce macrophage population in the uterus, but was rescued by Csf1 overexpression^52^. Tissue specific enhancement of Csf1 expression is accompanied with increased macrophage density^53^. Our results showing depletion HMGB1 with increased expression of *Csf1 mRNA* in deleted uteri suggest that Csf1 is required for macrophage retention. The regulation of local *Csf1* expression is unclear. Nonetheless, present evidence that HMGB1 is an important regulator of Csf1 induction and macrophage enrichment in the stroma.

Uterine receptivity to implantation is achieved through a cross-talk between the epithelium and stroma with specific gene expression in these two tissue types. It is interesting to note that epithelial receptivity marker genes *Ihh* and *Msx1* were comparable in floxed and deleted uteri, whereas stromal markers *Hoxa10* was substantially reduced. These defects were reflected in reduced expression of *Ptgs2* and *Bmp2*, two critical markers of blastocyst attachment and decidualization. These results suggest that uterine receptivity was not fully achieved in *Hmgb1*^d/d^ females. This is a unique observation that was revealed in this study.

Taken together, our study addresses a potential mechanism to control inflammation during early pregnancy and its effect on implantation. The unappreciated role of HMGB1 in balancing inflammation with respect to macrophage distribution depicts a new aspect for fertility and pregnancy events. Whether HMGB1 bears similar roles in other species including humans will require careful studies.

## Methods

### Mice

*Pgr^Cre/+^*, *Hmgb1*^*f/f*^ mouse lines were generated as previously described ^16,17^. *Hmgb1*^*d/d*^ mice were generated by mating floxed females with *Pgr*^*Cre/+*^ males. All mice used in this study were housed under a constant 12-h/12-h light/dark cycle in the Cincinnati Children’s Animal Care Facility according to NIH and institutional guidelines for the use of laboratory animals. All protocols were approved by the Cincinnati Children’s Animal Care and Use Committee. Mice were provided *ad libitum* with autoclaved Laboratory Rodent Diet 5010 (Purina) and UV light-sterilized reverse osmosis/deionized constant-circulation water ad libitum.

### Analysis of pregnancy events

Pregnancy events were assessed as previously described^3,4,5,6,44,45^. Three adult females were randomly chosen and housed with a WT fertile male overnight in separate cages; the morning of finding the presence of a vaginal plug was considered successful mating (day 1 of pregnancy). Plug-positive females were then housed separately from males until processed for experiments. Litter size, pregnancy rate, and outcomes were monitored in timed pregnancy. Blue reaction was performed by injecting intravenously a blue dye solution (1% Chicago Blue in Saline, 100 μL/mouse) 4 min prior to mice were killed. The distinct blue bands along the uterus indicated the site of implantation. For confirmation of pregnancy in plug-positive day 4 mice or mice showing no blue bands on day 5, one uterine horn was flushed with saline to confirm the presence of blastocysts. If blastocysts were present, the contralateral horn was used for experiments and mice without any blastocyst were discarded. Pseudopregnancy was induced by mating females with vasectomized males. For rescue experiments, progesterone (P4, 2mg/100μl/dose) was injected subcutaneously to pregnant mice on the morning of days 3 and 4. Mice were sacrificed after blue dye injection on day 5 of pregnancy. Tissues were frozen and proceeded to IF to examine macrophage accumulations.

### Histology

Tissue sections from control and experimental groups were processed on the same slide. Frozen sections (12 μm) were fixed in 4% PFA-PBS for 10 min at room temperature and then stained with hematoxylin and eosin for light microscopy analysis.

### Immunofluorescence (IF) and Microscopy

IF was performed as previously described^6,45^. IF using frozen sections (12 μm) was performed using the following 1^st^ antibodies: ERα (1:300, sc-542; Santa Cruz), PR (1:300, 8757; Cell Signaling technology), FOXA2 (1:300, WRAB-FOXA2, Seven Hills Bioreagents), E-Cadherin (1:300, 3195s, Cell Signaling Technology), CK8 (1:1000, TROMA-1, Hybridoma Bank, Iowa), Ki67 (1:300, RM-9106-S, Thermo Fisher Scientific), Cleaved Caspase-3 (1:300, 9661s, Cell Signaling technology), CD45 (1:300, 103012, Biolegend), F4/80 (1:1000, MCA497R, Biorad), MHC II (1:500, 14-5321-81, eBioscience), CD206 (1:1000, MCA2235T, AbD Serotec), CSF1R (1:1000, sc-692, Santa Cruz), Gr-1 (1:500, MCA2387, Bio-Rad), Fluorescein labeled DBA-lectin (1:500, FL-1031-2, Vectorlabs) and CD31 (1:300, BD 553370, BD Biosciences). For IF on paraffin sections (6 μm), anti-HMGB1 antibody (1:2000, 6893S, Cell Signaling Technology) and anti-CD68 antibody (1:1000, MCA1957T, AbD Serotec) were used. For signal detection, secondary antibodies conjugated with Cy-2, Cy-3, Alexa 488, or Alexa 594 (1:300, Jackson Immuno Research) were used. Nuclear staining was performed using Hoechst 33342 (5 μg/ml, H1399, Thermo Fisher Scientific). Tissue sections from control and experimental groups were processed onto the same slide. Pictures were taken using Nikon Eclipse 90i upright microscope and processed by Nikon Elements Viewer. Cell number counting was performed using Image J (NIH).

### In Situ Hybridization

Frozen sections from *Hmgb1*^*f/f*^ or *Hmgb1*^*d/d*^ mice were processed onto the same slide. *In situ* hybridization with ^35^S-labeled cRNA probes was performed as described^3,4^. Digoxigenin (DIG)-labeled probes were generated according to the manufacturer’s protocol (Roche). *In situ* hybridization with DIG-labeled probes was performed as described^44^. The primers used for the DIG-labeled probe of *Hmgb1* are listed below: 5’- CGGATGCTTCTGTCAACT-3’ and 5’- ACTTCTCCTTCAGCTTGG-3’.

### Cytoplasm/Nuclear protein fractionation

Pregnant uterine tissues were homogenized in Buffer B (5 mM EDTA-PBS). Tissue homogenates were centrifuged at 1,000g, 4 °C for 2 min and supernatants were discarded. Remaining pellets were resuspended in Buffer A (10 mM HEPES pH7.9, 10 mM KCl, 0.1 M EDTA, 1 mM DTT, 0.5 mM PMSF) and kept on ice for 20 min. After adding 1/4 volume of Buffer A with 2.5% NP-40 and vortex, samples were centrifuged at 15,000 g, 4 °C for 5 min. Supernatants were collected as cytoplasm fractions and pellets were vortexed in Buffer C (20 mM HEPES pH7.9, 0.4 M NaCl, 1 mM EDTA, 1 mM DTT, 1 mM PMSF) at 4 °C for 25 min. Supernatants were collected as nuclear fractions after centrifugation at 17,000 g, at 4 °C for 5 min. All samples were kept at −20 °C until use.

### Immunoblotting

Western blotting was performed as described^3,4^. The same blots were used for quantitative analysis of each protein. Bands were visualized using an ECL kit (Bio-Rad). α-Tubulin and Lamin A/C served as a loading control. The following antibodies were used to detect each protein: HMGB1 (1:1000, 6893S, Cell Signaling Technology), α-Tubulin (1:1000, 2144, Cell Signaling Technology) and Lamin A/C (1:1000, sc20681, Santa Cruz).

### Isolation of primary stromal cells

Stromal cells from day 4 of pregnant uteri were collected by enzymatic digestion as described previously^3^. Uteri from *Hmgb1*^*f/f*^ or *Hmgb1*^*d/d*^ mice on day 4 of pseudopregnancy were split open longitudinally and cut into small pieces (2-3 mm long). Tissue pieces were incubated with pancreatin (25 mg/mL, Sigma) and dispase (6 mg/mL, Gibco) for 1 h at 4 °C, followed by 1 h at room temperature and 15 min at 37 °C. Luminal epithelial (LE) sheets were removed by pipetting tissues several times. The remaining tissue fragments were incubated in type IV collagenase (300 U/mL, Washington) to free stromal cells. Stromal cells were suspended in DMEM: F12 Nutrient Mixture (Hyclone) containing 10% heat-inactivated FCS (Gibco), 50 units/mL penicillin, 50 μg/mL streptomycin and 1.25 μg/mL fungizone (Pen Strep; Gibco). Cell suspensions were filtered through a 70-μm nylon mesh to remove glands and clumps of epithelial cells. Cells were seeded into 6 well plates and medium was changed 1 h later to remove unattached immune cells. After another 5 h culture, cells were washed in PBS and dissolved in TRIzol reagent.

### RT-PCR

RT-PCR was performed as described^3,4^. PCR was run for 25 cycles using following primers: 5’- AGATGACAAGCAGCCCTAT-3’ and 5’- CTTTTCAGCCTTGACCAC-3’ for *Hmgb1*; 5’- GCAGATGTACCGCACTGAGATTC-3’ and 5’-ACCTTTGGGCTTACTCCATTGATA-3’ for *Rpl7*; Rpl7 served as an internal control. Each PCR product was loaded onto 2% Agarose gel containing ethidium bromide with a volume of 3μl to detect target bands.

### Quantitive RT-PCR

RT-qPCR was performed as described^3,4^, using the following primers: 5’-AGAAGCTGTAGTTTTTGTCACC-3’ and 5’- TGCTTGAGGTGGTTGTGGAA-3’ for *Ccl2*; 5’- CTCTAGCCGAGGCCATGTGGAG-3’ and 5’-GGCCCCCAACAGTCAGCAAG- 3’ for *Csf1*; 5’-GCAGATGTACCGCACTGAGATTC-3’ and 5’-ACCTTTGGGCTTACTCCATTGATA-3’ for *Rpl7*; *Rpl7* served as an internal control.

### Luciferase assay for PR and GR

Luciferase assay was performed as previously described^32^, using the primary cultured stromal cells. Briefly, 1 × 10^5^ cells were seeded in 24 well dishes. After 48 h, cells were transfected with 250 ng of PRE2-TK-Luc and 2 ng of pRL-TK by Lipofectamine 2000 (Invitrogen) according to the manufacturer’s protocol. Six hours later, media were changed to 10% FBS-containing DMEM/F12 and incubated for 18 h. Cells were then starved for 4 h, followed by the treatment of either medroxy-progesterone Acetate (1 μM) or dexamethasone (0.1 μM) for 20 h. Collected cells were proceeded to luciferase assay according to manufacturer’s protocol (Promega).

### Migration Assay

Before assay, Raw264.7 cells were cultured overnight in 1% FBS-DMEM. Cells were then suspended in serum-depleted DMEM at the concentration of 2.5 × 10^5^ cells/mL. In each bottom well, 700 μL of DMEM were added with or without 100 ng/mL Csf1 (R&D), 100 ng/mL Ccl2 (Biolegend), 1 μM Csf1r inhibitor (GW2580; Calbiochem) and 10 μM Ccr2 antagonist (BMS CCR2 22; Calbiochem). Upper inserts received 100 μL of DMEM with or without 3 μM Csf1r inhibitor or 30 μM Ccr2 antagonist and then 200 μL of cell suspensions. After culture of 4 h for Csf1 and 24 h for Ccl2, migrated cells were fixed in 4% PFA-PBS and stained in 0.1 % crystal violet for observations. Three fields/well at 8x magnification were quantified using ImageJ (NIH).

### Measurement of serum progesterone (P_4_) levels

Sera were collected on day 4 of pregnancy, and hormone levels were measured by enzyme immunoassay kits (Cayman) as previously described^3,4,5,6,44,45^.

### Tridimensional visualization of implantation sites

Whole mount staining, tissue clearing and 3D visualization of day 5 and 6 implantation sites were performed as previously described^6^. Anti-E-Cad antibody (1:100, 3195s, Cell Signaling Technology) and anti-F4/80 (1:1000, MCA497R, Biorad) were used to stain the lumen epithelium and macrophages, respectively, with Alexa 594 (1:250, Jackson Immuno Research) anti-rat antibody conjugated with Alexa 488 (1:250, Jackson Immuno Research) as second antibodies. 3D images were acquired by a Nikon multiphoton upright confocal microscope (Nikon A1R). To obtain the 3D structure of the tissue, the surface tool in Imaris (version 9.2.0., Bitplane) was used.

### Co-immunoprecipitation (Co-IP)

Co-IP was performed as previously described^6^. The following antibodies were used for Co-IP: GR (1:200, 12041S, Cell Signaling technology) and PR (1:200, 8757; Cell Signaling technology).

### *Hmgs*, *Csf1* and *Ccl2* expression status examined by RNA-Seq analysis

Whole uterine tissues or enzymatic digested epithelial/stromal cells on day 4 were homogenized in TRIzol to extract total mRNA. After removing genomic DNA, total RNAs were subjected to RNA-Seq by HiSeq2500. The expression levels of genes were presented as reads per kilobase per million (RPKM) by Tophat2^54^ and visualized by R.

### *Csf1*, *Ccl2* and *Csf1r* expression in different cells types in uterus by single cell sequencing

The database of whole uterus single cells sequencing from mouse cell atlas by microwell technology was applied. Seurat was applied to re-analysis expression of *Csf1*, *Ccl2* and *Csf1r* expression in different cell types of uterus (GEO: GSE108097)^37^.

### Statistics

Statistical analyses were performed using a two-tailed Student’s *t*-test or a multiway analysis of variance (ANOVA) using Prism 6 (GraphPad Software). A value of P < 0.05 was considered statistically significant.

## Acknowledgements

The authors really appreciate Katie Gerhardt for editing the manuscript. *Pgr-Cre* mice were originally obtained from Francesco DeMayo and John P Lydon (Baylor College of Medicine). Robert F Schwabe (Columbia University) originally provided the floxed *Hmgb1* mouse line. The vector for PRE2-Tk-Luc was gifted from Dean P Edwards (Baylor College of Medicine). This work was supported in part by NIH grants (HD068524 and DA006668) and March of Dimes Center grant (22-FY17-889) to S.K.D.. S.A. was supported by Astellas Foundation for Research on Metabolic Disorder Fellowship for Study Abroad and is now supported by Osamu Hayaishi Memorial Foundation Fellowship for Study Abroad.

## Author contributions

S.A., W.D., X.L., Y.J., A.B. and X.S. performed experiments, S.A., W.D., and S.K.D. designed experiments. S.A., W.D., X.S. and S.K.D. analyzed data. S.A., W.D. and S.K.D wrote the manuscript.

## Data availability

The authors declare that all data supporting the findings of this study are available within the article and its Supplementary Information files or from the corresponding authors on reasonable request. The sequencing data generated in this study have been deposited in the NCBI Gene Expression Omnibus (GEO) repository under the accession number GEO: GSE120549 and GSE116096

## Author Information

The authors declare no competing financial interests.

---

**Supplementary Figure 1.**
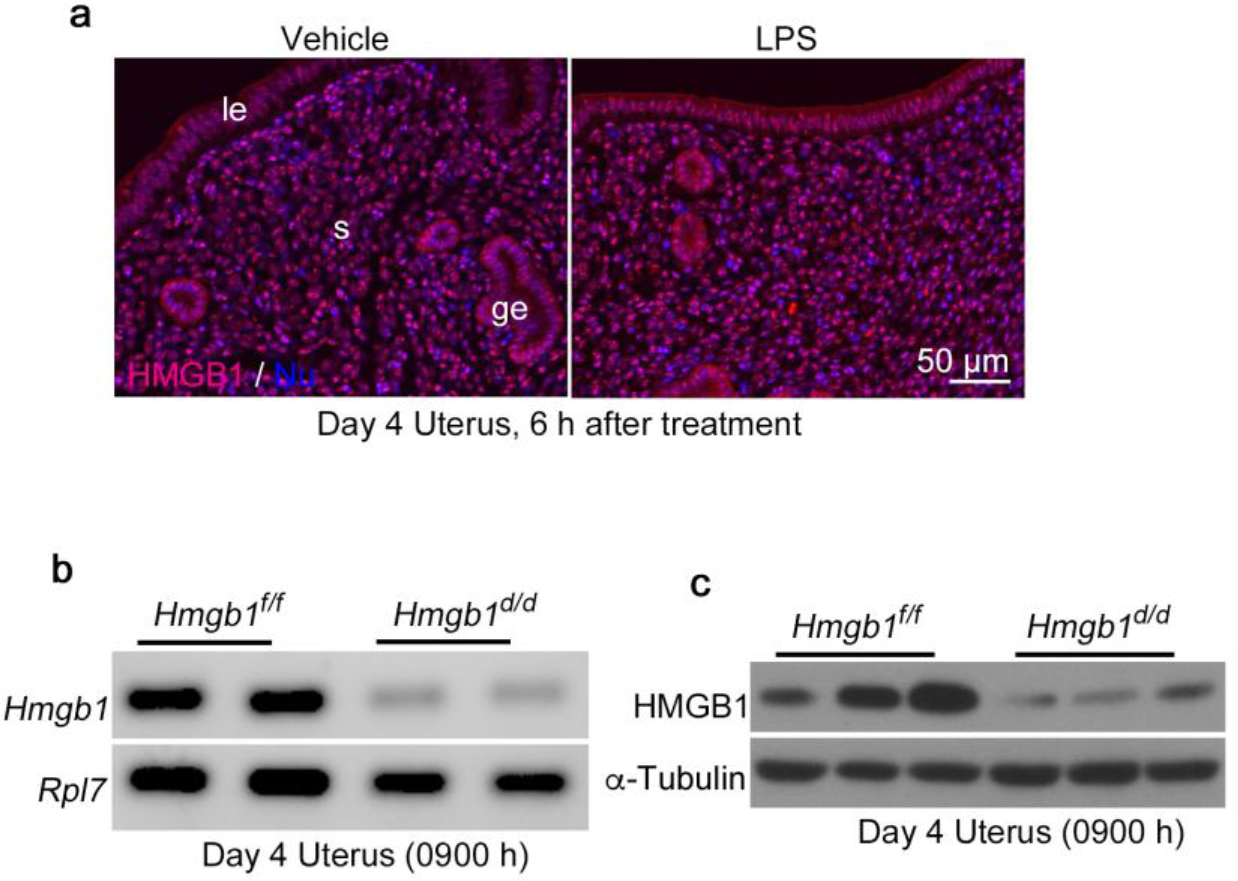
Expression of HMGB1 in Pregnant Uteri. **a**, IF of HMGB1 in day 4 uteri of *Hmgb1*^*f/f*^ mice 6 h after i.p. injection of 2.5 μg/mouse LPS. le: luminal epithelium, ge: glandular epithelium, s: stroma. Scale bar: 50 μm. Each image is a representative from at least 3 independent experiments. **b**, **c**, Deletion efficiency of *Hmgb1* in day 4 uteri as examined by RT-PCR and Western-blot. *Rpl7* and α-Tubulin were used as loading controls.

**Supplementary Figure 2.**
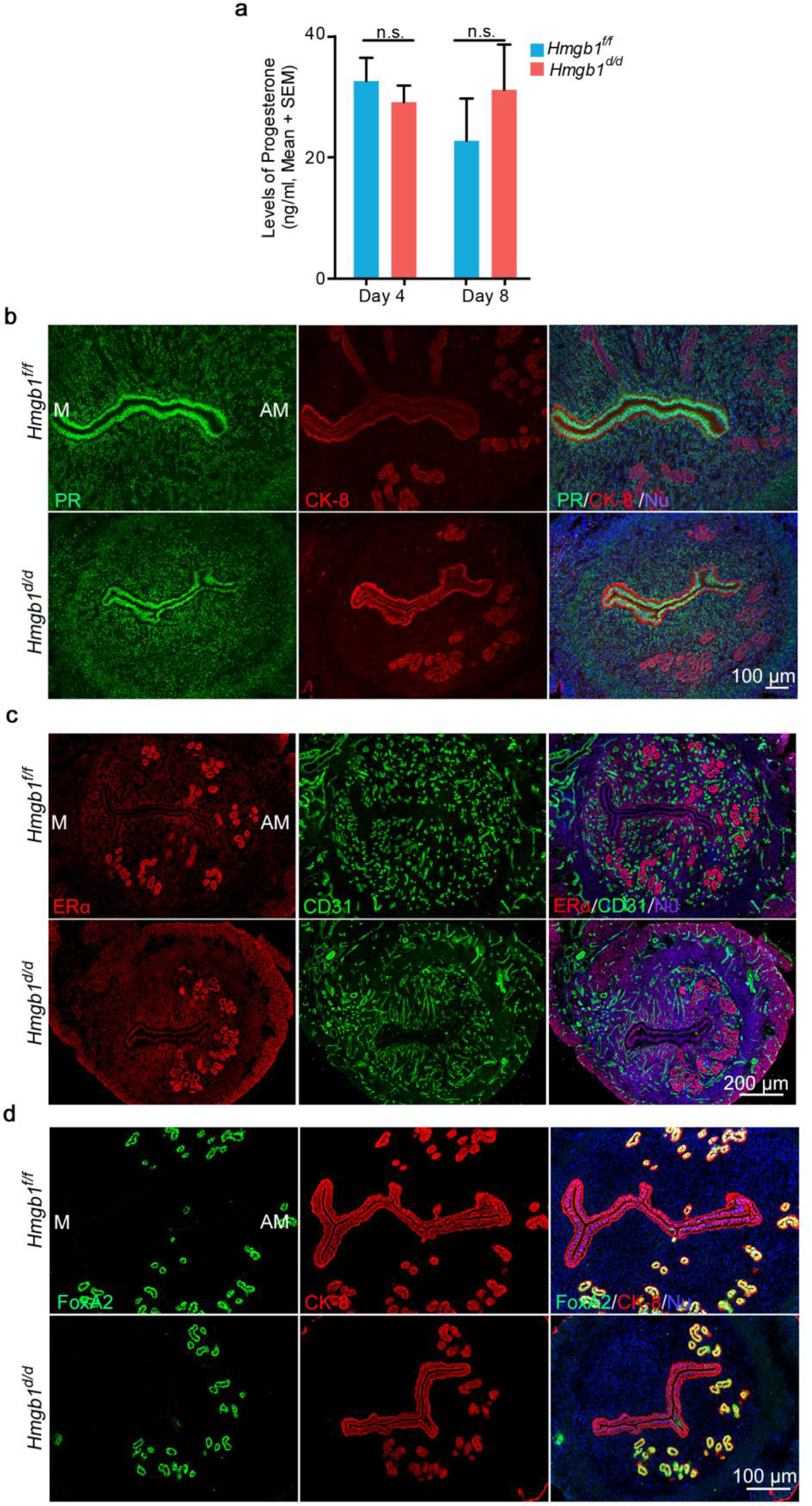
Serum P_4_ Levels and Status of PR, ERα and FoxA2 are comparable between floxed and deleted uteri. **a**, Serum P_4_ levels on days 4 and 8 were analyzed by Progesterone EIA kit. n = 5 for each genotype. Data are presented as mean ± SEM, n.s.: not significant by student’s *t*-test. **b**, IF of PR and CK8 in day 4 pregnant uteri from *Hmgb1*^*f/f*^ and *Hmgb1*^*d/d*^ females. M: mesometrial pole, AM: antimesometrial pole. Scale bar: 100 μm. **c**, IF of ERα and CD31 in day 4 pregnant uteri from *Hmgb1*^*f/f*^ and *Hmgb1*^*d/d*^ females. M: mesometrial pole, AM: antimesometrial pole. Scale bar: 200 μm. **d**, IF of FoxA2 and CK8 in day 4 pregnant uteri from *Hmgb1*^*f/f*^ and *Hmgb1*^*d/d*^ females. M: mesometrial pole, AM: antimesometrial pole. Scale bar: 100 μm. Each image is a representative from at least 3 independent experiments.

**Supplementary Figure 3.**
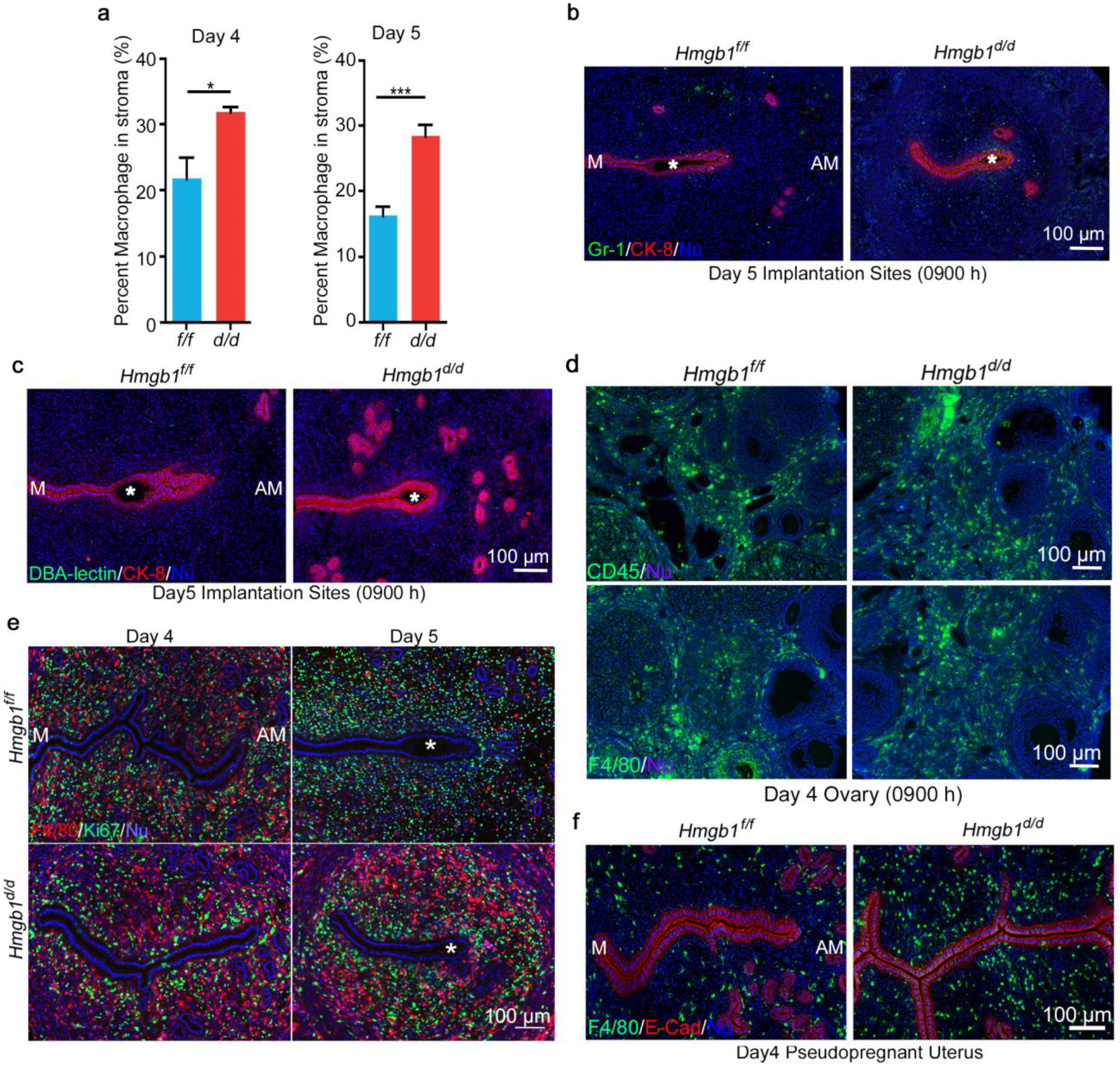
*Hmgb1* Deletion in Uteri Does Not Affect to Neutrophils and NK Cells in Uteri and Leukocytes in Ovary. **a**, Number of F4/80 positive cells in the stromal layer per section in **Figure 4b** was counted by Image J software. Data are presented as mean ± SEM, n = 5 for each genotype in day 4 and n = 6 for each genotype in day 5. \**P* < 0.05 and \*\*\**P* < 0.001 by student’s t-test. **b,** IF of Gr-1 (neutrophil marker) and CK8 in day 5 implantation sites from *Hmgb1*^*f/f*^ and *Hmgb1*^*d/d*^ females. Asterisks indicate blastocysts. M: mesometrial pole, AM: antimesometrial pole. Scale bar: 100 μm. **c,** IF of DBA-lectin (NK cell marker) and CK8 in day 5 implantation sites from *Hmgb1*^*f/f*^ and *Hmgb1*^*d/d*^ females. Note absence of NK cells in stroma of both genotypes. Asterisks indicate blastocysts. M: mesometrial pole, AM: antimesometrial pole. Scale bar: 100 μm. **d,** IF of CD45 and F4/80 in day 4 ovaries from *Hmgb1*^*f/f*^ and *Hmgb1*^*d/d*^ females. Scale bar: 100 μm. **e**, IF of F4/80 and Ki67 in days 4 and 5 pregnant uteri from *Hmgb1*^*f/f*^ and *Hmgb1*^*d/d*^ females. Asterisks indicate blastocysts. M: mesometrial pole, AM: antimesometrial pole. Scale bar: 100 μm. Each image is a representative from at least 3 independent experiments. **f,** IF of F4/80 and E-Cad in day 4 pseudopregnant uteri from *Hmgb1*^*f/f*^ and *Hmgb1*^*d/d*^ females. M: mesometrial pole, AM: antimesometrial pole. Scale bar: 100 μm.

**Supplementary Figure 4.**
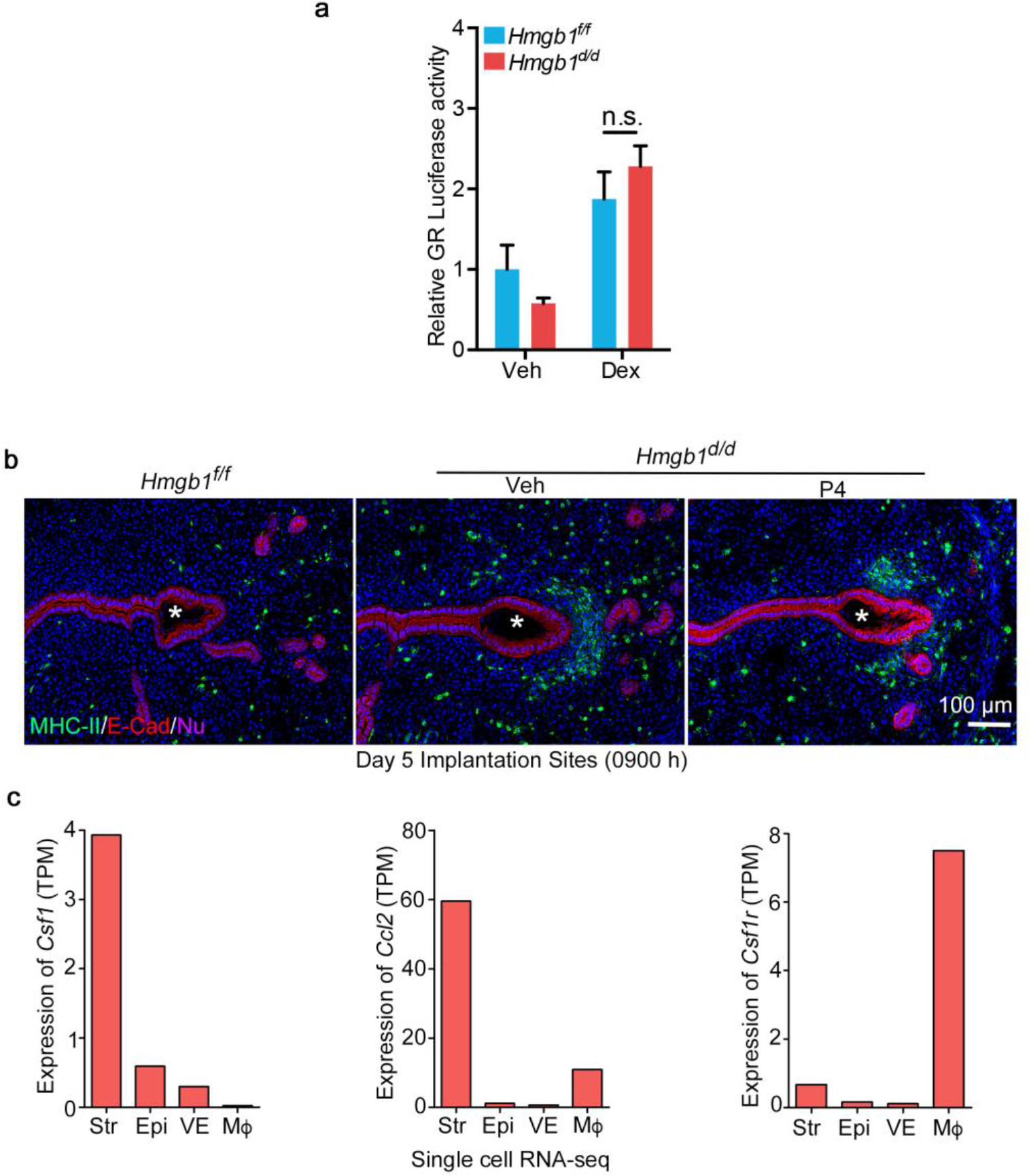
HMGB1 Regulates Macrophage Migration via Suppression of Macrophage Attractants. **a**, Luciferase assay shows GR activity is comparable between *Hmgb1*^*f/f*^ and *Hmgb1*^*d/d*^ stromal cells. n = 3 for each genotype. Data are presented as mean ± SEM, n.s.: not significant by One-Way ANOVA. **b**, IF of MHC-II and E-Cad in day 5 implantation sites from *Hmgb1*^*f/f*^ and *Hmgb1*^*d/d*^ females (control or P_4_ treated). Asterisks indicate blastocysts. Scale bar: 100 μm. Each image is a representative from at least 3 independent experiments. **c**, Expression levels of macrophage attractants, *Csf1* and *Ccl2*, and *Csf1r* were analyzed by single cell RNA-seq in non-pregnant uteri. Str: Stroma, Epi: Epithelium, VE: Vascular Endothelium, Mϕ: Macrophage.

**Video S1: Movie of 3D imaging of a Day 5 implantation site in *Hmgb1*^*f/f*^ mice (related to Fig. 4c).**

**Video S2: Movie of 3D imaging of a Day 5 implantation site in *Pgr*^*Cre/+*^*Hmgb1*^*f/f*^ mice (related to Fig. 4c).**

